# Laboratory *in vitro* replication of Ostreid Herpes Virus (OsHV-1) using Pacific oyster tissue explants

**DOI:** 10.1101/2024.05.10.593497

**Authors:** Robert W.A. Potts, Tim Regan, Stuart Ross, Kelly Bateman, Chantelle Hooper, Richard Paley, Ross D. Houston, Tim P. Bean

**Affiliations:** The Roslin Institute and Royal (Dick) School of Veterinary Studies, University of Edinburgh, Edinburgh, United Kingdom; Centre for Environment Fisheries and Aquaculture Science (Cefas) Weymouth Laboratory, Dorset, United Kingdom; Centre for Sustainable Aquaculture Futures, University of Exeter, Exeter, United Kingdom

## Abstract

Pacific oysters (*Crassostrea* or *Magallana gigas*) are one of the most economically important aquaculture species globally. Over the past two decades, ostreid herpesvirus (OsHV-1), has become a major pathogen of cultured Pacific oysters resulting in widespread mortality with a global distribution. Experimental use of OsHV-1 is challenging for many reasons, including both complexity and relative obscurity of host pathogen dynamics, and a lack of functioning model systems. The goal of this study was to improve the tools available for working with OsHV-1 in both whole animals and in tissue explants established *ex vivo* from oysters and maintained in controlled laboratory conditions. Tissue explants were taken from oysters originating from two different sources that have different levels of mortality in OsHV-1 challenges and were used in disease challenges alongside whole animals for comparison. Quantitative PCR, histology and electron microscopy were used to confirm that the explants were capable of replicating OsHV-1. Furthermore, the quantitative PCR results suggests that the source of the oysters was significant in determining the outcome of infection in the explants, supporting the validity of the explant model for OsHV-1 infection. This approach for studying OsHV-1 allows for the control of confounding factors in disease outcome that is not possible in whole animal challenges, providing a new tool for studying OsHV-1 in Pacific oysters.

## Introduction

Ostreid Herpesvirus 1 (OsHV-1) is an important pathogen of Pacific oyster aquaculture and has been the subject of studies worldwide. However, systems for studying host pathogen interactions of OsHV-1 and Pacific oysters are limited. Experiments have previously been conducted in the open ocean where they rely upon natural outbreaks and where there are a wide range of factors that can potentially influence the outcome of infection (King et al., 2019). For example, abiotic factors can play an important role in infection by stressing the host, which can thus reduce ability to prevent or tolerate infection or create conditions ideal for pathogen proliferation (Pernet et al., 2014). Younger oysters are most prone to OsHV-1 infection (Dégremont et al., 2015, Azema et al., 2017), and are also more susceptible to changes in salinity and temperature (Nell and Holliday, 1988, Malham et al., 2009). These changes can be difficult to control in open environments or hatcheries, and are usually a function of normal ecosystem variation.

A greater level of control is achieved by infecting whole oysters with OsHV-1 in systems, such as recirculating aquaria or even in static aquaria (Hick et al., 2018, Friedman et al., 2020, Burge et al., 2020). Abiotic factors such as temperature and salinity can be tightly restricted using incubators and artificial or purified seawater (Burge et al., 2020). However, it is not possible to completely control for variability. For example, animals are commonly exposed to the naturally occurring microbiota prior to controlled disease challenges as there are few options for production of truly axenic whole animals. The interaction between OsHV-1 and facultative pathogens, such as *Vibrio spp.* bacteria, is complex and can play a major role in the outcome of infection (de Lorgeril et al., 2018, Fallet et al., 2022). Furthermore, the microbiome of the oyster plays a key role in health and disease susceptibility of the host (King et al., 2019) and there remains a vast diversity of marine microorganisms that have not been studied which may interact with the aetiology of oyster diseases (Hartmann et al., 2014, King et al., 2019). Detecting and removing microorganisms from experimental animals, and thus from recirculating or static environments, can be challenging. Also, the experimental environment often provides ideal conditions for proliferation, further affecting the desired experimental conditions.

A further issue with using whole animals is inconsistency between individuals. A key example of this is behavioural e.g. oysters infected using a bath challenge method may not open and filter the surrounding water in the time between initiation of the challenge and the degradation of the OsHV-1 viability (thought to be within 24 hrs within experimental systems), resulting in unsuccessful models of infection. Injecting OsHV-1 directly into the adductor muscle has been used to guarantee ingress of a known quantity of virus, but this is less relevant to a natural infections where exposure will occur through gill and/or feeding apparatus. Hemocytes are another potential model for OsHV-1 infection, as they play a key role in host response to infection and have been shown to facilitate OsHV-1 replication *in vivo* (Morga et al., 2017). However, hemocyte cultures from Pacific oysters do not proliferate and are short lived, making them difficult to work with for experiments lasting over 24 hours, which are crucial for studies of complete viral life cycles (Li et al., 2015). Hemocytes from the scallop *Chlamys farreri* have been maintained for long enough to show that OsHV-1 can successfully replicate in hemocyte culture, although this has not been applied to *Crassostrea gigas* (Ji et al., 2017). Attempts to replicate OsHV-1 in primary cell types other than hemocytes has so-far proved unsuccessful (Renault et al., 2002).

Creating a model that is representative of the disease dynamics that occur in aquaculture systems whilst removing as many confounding factors as possible would be a major addition to the OsHV-1 research toolkit. The ability to control factors such as temperature, pH, microbiology, and salinity in a laboratory setting would allow for better identification of causative factors in the complexity of disease aetiology. The improvement in primary cell cultures in Pacific oyster offers one potential option (Potts et al., 2020). Cell culture systems lend themselves to controlling abiotic conditions through buffering of pH and temperature, as well as simplified and standardised materials. Disease challenges in enclosed, controlled cell culture environments reduces both the technical, logistical and biosecurity challenges associated with disease challenges in aquaria or natural environments. The ability to exclude or introduce different biotic factors such as *Vibrio* bacteria would help to elucidate the complex relationships between these organisms and outcomes of OsHV-1 infection. Oysters can be reared in axenic conditions, but this requires starting at the larval stage (Douillet and Langdon, 1993). Removing the microbiota from the oyster without too much stress is a challenge that may be overcome with a cell culture approach. With any model system it is essential to assess the benefits of simplification against any loss of biological relevance to the field situation: This can partially be done by reassessing the difference in outcomes between the whole animals and the cell culture model. If successful, this approach could then be applied to different molluscan aquaculture species and the wide range of diseases that impact them (Potts et al., 2021).

Here, we demonstrate the effective application of *ex vivo C. gigas* culture systems of several different tissues to study OsHV-1 µvar infection in two different populations of Pacific oyster. The *ex vivo* model enabled us to quantify the level of viral replication in the different tissue types. Use of both *in vivo* and *ex vivo* challenge models in two oyster populations with different susceptibility to OsHV-1 provided a comparison of the two challenge methods to validate the use of the tissue explant model for OsHV-1 infection.

## Materials and Methods

### Preparation of OsHV-1 Stock

OsHV-1 was acquired from a preserved master stock “MS8” (approximately 3×10^5^ viral copies μl^-1^) produced by The Centre for Environment, Fisheries and Aquaculture Science (Cefas) laboratory in Weymouth, derived originally from a natural outbreak of OsHV-1 in Poole Harbour in 2015 (see Morga et al., 2021 for viral genome analysis). To amplify viral stocks, oysters approximately 20 mm in height (umbo to bill) were anaesthetised in aerated magnesium chloride seawater solution (50 g/L) for 4 hours (Suquet et al., 2009). Oysters with open valves were split into four groups of 12, transferred to 6 well plates and the adductor muscle injected with a 40-gauge needle. The first group was injected with 50 µl OsHV-1 MS8 stock immediately after thawing, the second group with a 1 in 10 dilution of OsHV-1 MS8 stock, the third group with a 1 in 100 dilution of MS8 OsHV-1 stock and the final group was injected with 0.22 µm filtered artificial seawater at 35 ppt (ASW) as a control. Each well was filled with approximately 10 ml of filtered ASW, to ensure the oyster is completely covered with water. Plates were then incubated at 20°C in a dark incubator. Oysters were checked daily for mortality by tapping the shell with forceps, any animals that remained open were assumed dead or moribund. Water was replaced every 48 hours. Moribund or dead oysters were bagged and transferred to 4°C until the end of the challenge. At the end of the challenge, gill and mantle was dissected out, pooled, then homogenised using Stuart SHM10 tissue homogenizer and disposable probes. Homogenate was then diluted in ASW (1:1 volume) and filtered in series though 5 µm, 1 µm, 0.45 µm and 0.22 µm syringe filters. Aliquots of virus were prepared for all subsequent challenges (labelled as master stock 9; MS9). Aliquots of virus and homogenate control (prepared in the same way but from uninfected oysters) were also mixed with Glycerol (10% final volume) as a cryoprotectant.

Control animals were held in separate 6 well plates. Live animals were dissected in the same way as dead challenged animals at the end of the challenge. Homogenate was treated in the same way as infected stock and stored as homogenate control for subsequent challenges.

### Whole animal disease challenge

Juvenile oysters (approximately 20 mm in height - exact age unknown) were acquired directly from nursery systems of two different commercial oyster hatcheries in the UK. Oysters were maintained in a biosecure oyster holding facility until use. Small oysters were kept in 15 l Polypropylene tanks in groups of approximately 100 oysters in ASW. Tanks were aerated and oysters were fed algal paste diet of shellfish diet 1800 (Reed Mariculture) according to manufacturers’ recommendation, and water was replaced every two days. Oysters from different sites were held in identical conditions but in separate tanks.

Whole animal infection challenges were done to ensure the viability of the cryopreserved OsHV-1 masterstock 9 (OsHV-1 MS9). Animals were placed in 6 well plates that were filled with 10 ml ASW. OsHV-1 MS9 aliquots were thawed on ice and 100 µl was immediately added to each well. Oysters were checked for mortality daily and water was replaced every 48 hours. Dead oysters were stored at -20°C before being disposed of according to Roslin Institute aquaculture biosecurity measures plan.

To compare lethality of OsHV-1 in different populations, 30 animals were subjected to a bath challenge with 100 µl OsHV MS9 (2×10^4^ copies per µl), 30 animals were incubated with 10 µl OsHV-1 MS9, 30 animals were incubated with homogenate control material, and 30 animals were incubated in filtered ASW only. Each well was filled with ASW, so that each oyster was in approximately 10 ml of ASW before virus was added. This process was duplicated for each of the two oyster population sources.

### Explant infection

Explants for infection experiments were prepared from juvenile oysters from the two different commercial hatcheries. Whole or partial gill, mantle and adductor muscle tissues were removed from the oysters. For mantle and gill tissues, as much tissue as possible was dissected in a single piece whilst ensuring no contamination from other tissue types. Samples that were particularly small or potentially contaminated with other tissues were not used for the challenge. Samples were randomly distributed into challenge or control groups for each oyster source. Explants were held in 1 ml oyster media (50:50 ASW:L15 – see Potts et al., 2020) in 24 well plates for 24 hours prior to inoculation with virus. OsHV-1 MS9 (2×10^4^ copies per µl) was thawed by a short incubation at 20°C before 10 µl was immediately added to each infection well.

### Tissue fixation and OsHV quantification

Two small pieces of tissue (1 to 2 mm^3^) were cut from each tissue type and stored in 2% glutaraldehyde in cacodylate buffer for electron microscopy. Remaining tissue was immersed in RNAlater (Invitrogen), snap frozen and stored at -80 °C prior to DNA extraction by Qiagen DNeasy blood and tissue kit according to manufacturer’s protocol. Eluted DNA was also used as a template for qPCR, as above. The absolute number of viral copies per µl was calculated from Ct values using a standard curve generated using a known quantity of plasmid containing a cloned target sequence.

### Media OsHV-1 quantification

10 µl of media was taken from directly above the tissue explant daily from each well and snap frozen. Samples were stored at -20°C. 1 µl was used directly as template for OsHV-1 quantification via qPCR. qPCR was performed using ABI FAST 7500 real time PCR machine using FAM based probe detection in 20 µl reaction volumes, 10 µl of TaqMan Fast Advanced Master Mix for qPCR with 1/20,000 dilution of ROX reference dye, 0.5 µl primer OsHV BF (GTC GCA TCT TTG GAT TTA ACA A), 0.5 µl primer OsHV B4 (ACT GGG ATC CGA ACT GAC AAC), 0.5 µl OsHV-1 probe (FAM- TGC CCC TGT CAT CTT GAG GTA TAG ACA A -

TAM) (adapted from Martenot et al., 2010), 7.5 µl nuclease-free water and 1 µl media sample were analysed in 96 well plates. Thermal cycling programme consisted of 3 minutes at 95°C followed by 40 cycles of 95°C for 3 seconds and 60°C for 12 seconds. Ct values were used to calculate viral copies per µl using a standard curve generated from OsHV-1 amplicon inserted into a plasmid and aliquoted at a known concentration.

### Degradation of OsHV-1 DNA and viability

To test attenuation of viral infectivity in seawater, filtered ASW containing 4×10^3^ copies of virus μl^-1^ was incubated in well plates for 24 hrs or 48 hrs at 18°C in 6 well plates the dark before the addition of oysters (each treatment was replicated 18 times for a total of n=72 oysters, including a no-virus control). Mortality was assessed daily and compared to mortality of oysters added to 6 well plates containing the virus immediately after thawing of the OsHV-1 masterstock. This challenge was performed separately to those above but using identical husbandry conditions.

Four different mediums were used to test OsHV-1 viral DNA degradation over time in a 24 well plate. 1 ml of ASW, filtered seawater, reverse osmosis (RO) water or oyster media were aliquoted into 6 wells each for six replicates per medium. 1 µl of OsHV-1 MS9 was added to each well and mixed gently by pipetting. 1 µl was taken from the centre of each well and pooled by medium in a microcentrifuge tube. Samples were snap frozen and stored at -20°C before quantification. All samples were thawed and analysed simultaneously. 1 µl was taken from each pooled sample to use as template for quantitative PCR following steps outlined above.

### Transmission Electron Microscopy

Transmission Electron microscopy (TEM) was used to confirm OsHV-1 infection in tissue explants. Samples for TEM were selected based on qPCR quantification data, with samples most likely to be infected being chosen for further assessment, alongside control samples. Quality control and preliminary histology screening was completed using semi-thin sections.

Samples were removed from glutaraldehyde and washed twice in 0.1M Sodium cacodylate buffer for 15 minutes before post fixation in 1% osmium tetroxide for 1 hour. Tissues were rinsed in 0.1 M sodium cacodylate buffer and stored in cacodylate buffer overnight. Samples were dehydrated in a graded acetone series (10, 30, 50, 70, 90, 100, 100%) and embedded within Agar 100 resin (infiltrated with 1:2 then 2:1 mixtures of Agar 100 resin: acetone1, and finally 100% Agar 100 resin). Embedded samples were cut as semi thin sections (1-2 µm) and stained on glass slides with 1% Toluidine Blue solution. Slides were examined under 400x power light microscope. Samples with characteristic signs of OsHV-1 infection were selected for ultrathin sectioning. Between 3-5 ultrathin sections (70-90 nm) were cut from selected samples and mounted on copper grids. Grids were stained with aqueous uranyl acetate and Reynolds lead citrate (Reynolds, 1963). Sections were examined with a JEOL JEM 1400 transmission electron microscope, and digital images captured using an AMT XR80 camera and AMTv602 software.

### Statistical analysis

Analysis of survival data was done using Kaplan Meier survival analysis in GraphPad version 9.3.1. Statistical analysis of qPCR data was completed using MiniTab version 20.3. generalised linear models with pairwise analysis by Tukey’s test applied to media viral load and tissue viral load data.

## Results

### Attenuation of viral infectivity in sea water

OsHV-1 showed reduced virulence over time in seawater dropping from 66% survival when added directly alongside oysters, to 94% survival when oysters were added 24 hours after virus, and 100% survival following a 48 hour interval (Figure 1). No mortalities were observed in sea water controls over the 14 day challenge (data not shown).

**Figure 1.**
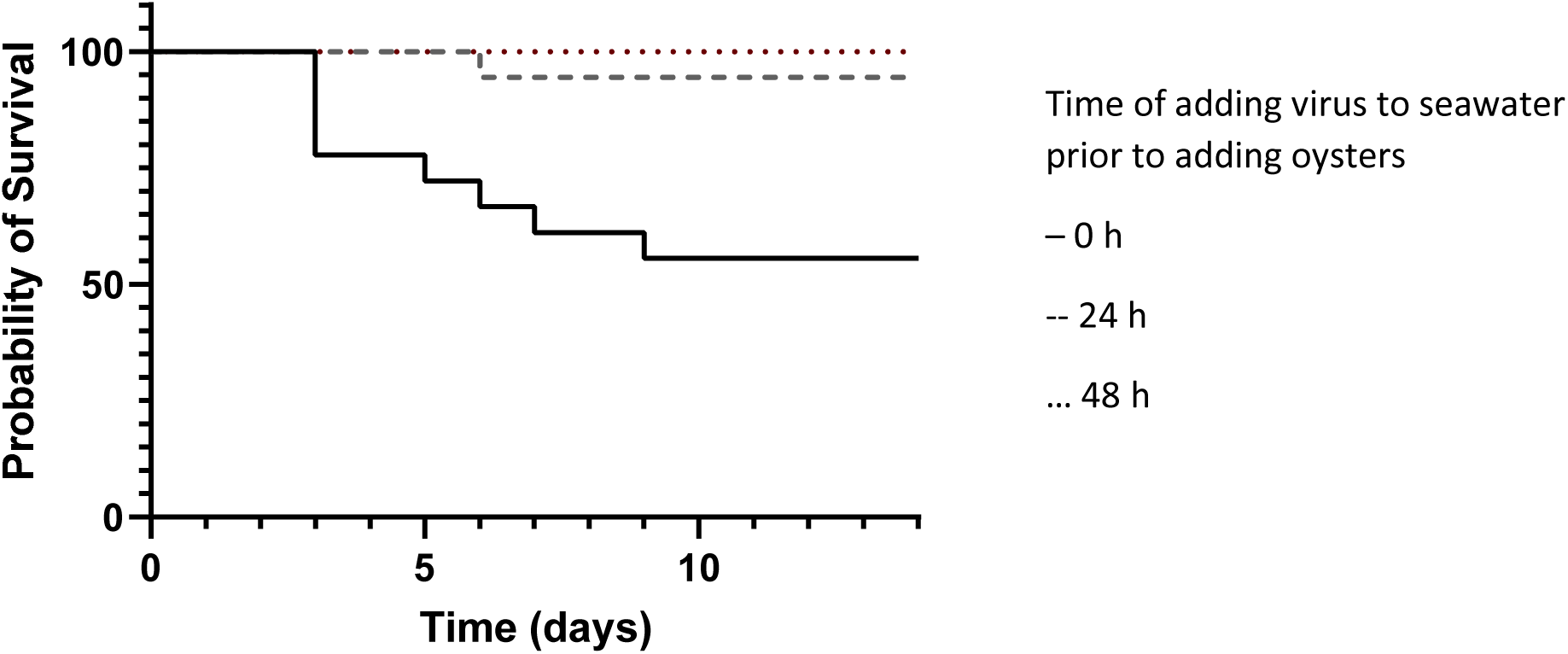
Attenuation of virus driven mortality in sea water over time. Virus was incubated for 0 (-), 24 (--), or 48 (···) hours prior to addition of oysters (n=18 per treatment) and mortality measured in oysters from this point onwards.

### Viral incubation in water

OsHV-1 detection by qPCR was consistent at around 30 cycles for ASW and oyster media over the 10 day experiment (Figure 2). Ct values increased slightly over time in RO water, from 29.1 to 34.0, suggesting a decrease in detectable viral DNA. Detection in natural raw seawater showed the greatest decrease, with Ct values rising from 31.3 on day 0 to 37.0 on day 8, after which no OsHV-1 DNA was amplified from the seawater samples.

**Figure 2.**
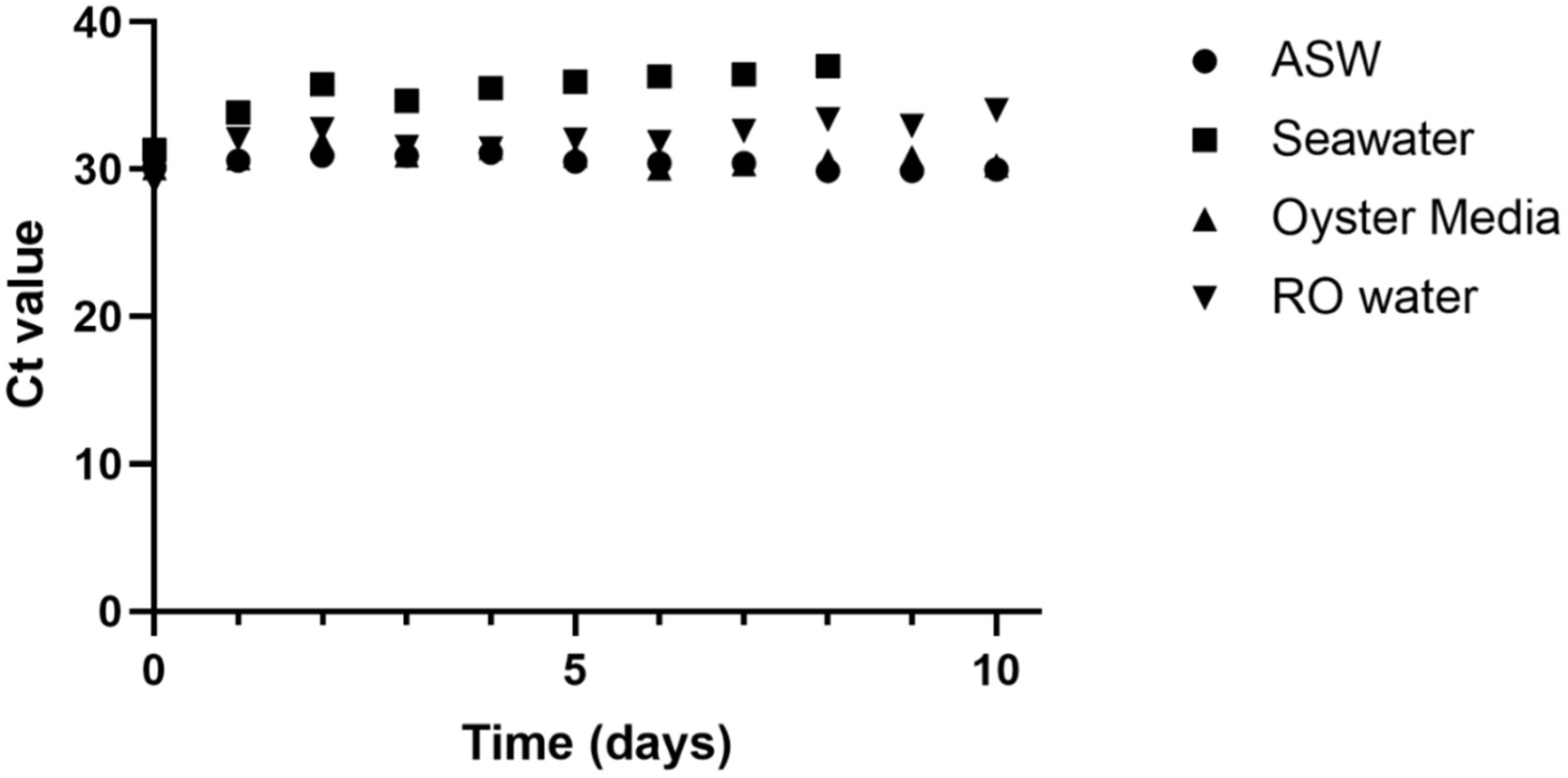
Ct values for viral DNA incubated in ASW, natural seawater, oyster media and RO water

### Comparison of mortality from different oyster sources

Animals from both sites in the seawater control group had 100% survival after the end of the challenge (Figure 3). Survival of animals incubated with oyster homogenate (without OsHV-1) was not significantly different to control (100% & 97% for site 1 and 2). Following viral insult, site one and site two had significantly different survival (Fisher’s exact test p<0.001) with site 1 having the lowest survival then treated with high dose (23%), followed by site 1 low dose (60%), site 2 high dose (77%) and site 2 low dose (87%).

**Figure 3.**
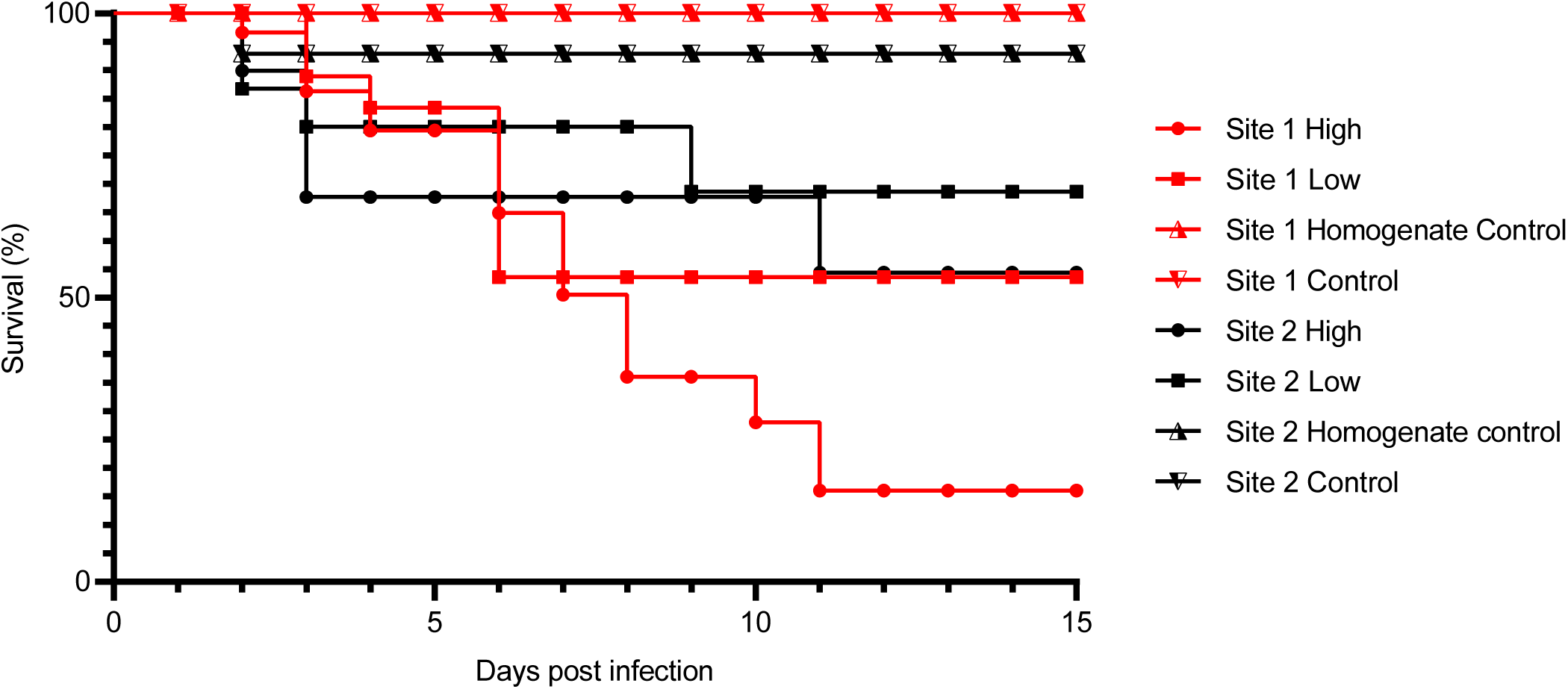
Survival of juvenile oysters from two different sites subjected to bath challenge with OsHV-1.

### OsHV-1 replication in tissue explant cultures

OsHV-1 quantification of DNA extractions from tissue explants ranged from below limit of detection to a maximum of 3.1×10^7^ copies μl^-1^of extract; approximately 100 times greater than the total amount of virus added to each well (2.4×10^5^). This amount was detected in DNA from a gill sample at 6 days post infection (Figure 4). As virus is unable to replicate in media alone (Figure 2), this further demonstrates that OsHV-1 is able to replicate within the tissue explant system developed. A generalised linear model with Tukey’s pairwise comparison showed viral load across all day was statistically significant for source of the oysters (p<0.001), driven mainly by significant differences between mantle and gill at each time point at each site. Levels of virus in tissues within each site were not significantly different. Over time there were various differences. Day 6 samples from site 1 mantle had significantly more virus than all day 2 and day 10 samples with the exception of site 1 day 10 gill and mantle, which were intermediate in viral load.

**Figure 4.**
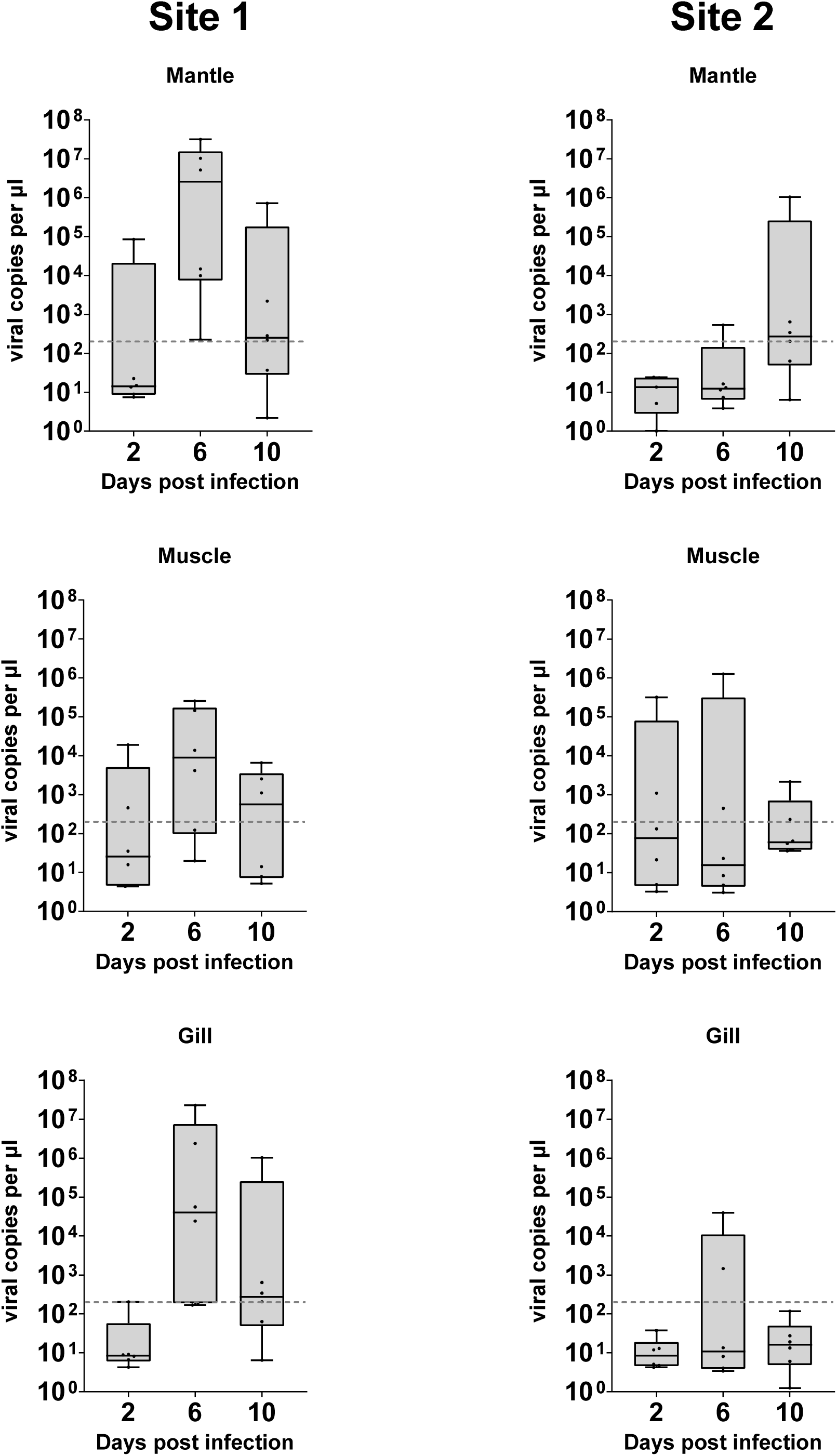
Virus copy number from DNA extraction of tissue explants challenged with OsHV-1 over time. Dotted line () represents initial viral copy number inoculated into media.

The presence of samples around or below the limit of detection suggests that infection was not present in all samples from any group, as with analysis of media. The frequency at which samples were well above this level was more frequent in groups from site one, and at later time points, but these groups are not significantly distinct.

OsHV-1 quantification of media surrounding oyster explants ranged from no detection to a maximum of 2.7×10^4^ copies μl^-1^, which was detected in media surrounding a gill sample from site 1 at 6 days post inoculation (Figure 5). A generalised linear model with Tukey’s pairwise comparisons of viral load of media samples demonstrated a statistically significant difference for source of the oysters (p<0.001), but with no significant difference of days post infection (p=0.076), or for different tissues. The only other statistically distinct group was the day 6 gill group from site one (p= 0.013), which had a significantly higher level of OsHV-1 DNA detected compared to other groups in the experiment.

**Figure 5.**
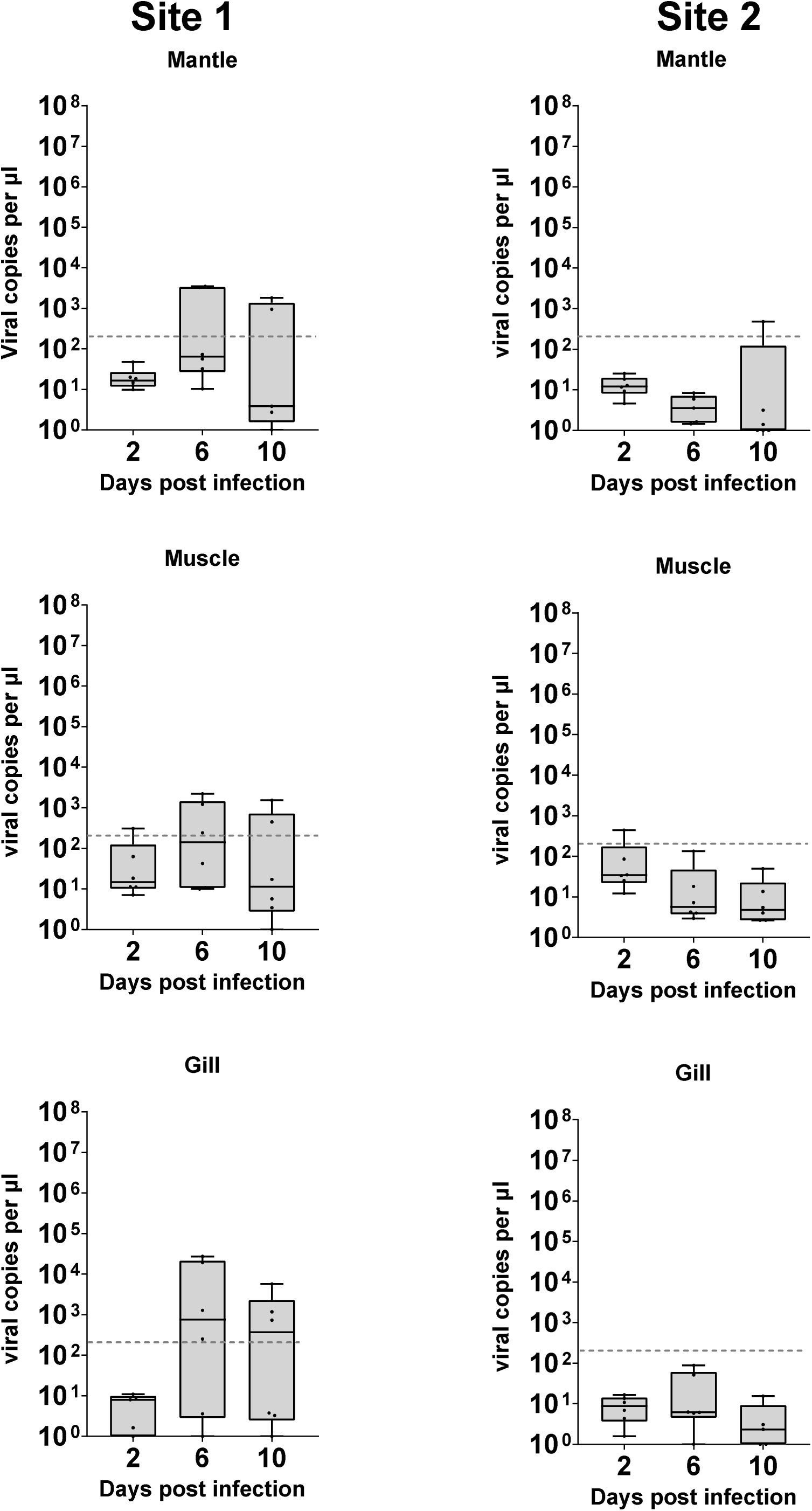
Virus copy number in media surrounding tissues challenged with OsHV-1 over time. Dotted line () represents initial viral copy number inoculated into media.

Across all groups there were samples that had quantifications less than 10 viral copies per µl, suggesting that infection was not present in all samples from any group. The frequency at which samples were well above this level was more frequent in groups from site one, and at later time points. Greater numbers of explants would need to be used to characterise the source of this variability.

### Quantification in tissue vs media

OsHV-1 was measured in the media itself at days 2, 6 and 10. Media Ct values ranged from a high of 21.7 to a low of 38.7, with the majority falling between 30 and 35 (Figure 5). The Ct values for detection from DNA extractions taken from tissue samples varied from 12.9 to 35.5, with a cluster in the 30 to 35 range. There is a correlation between the Ct values from the media and tissue for each sample. OsHV-1 detection in the DNA extractions from tissue explants was higher overall than in the media.

**Figure 5.**
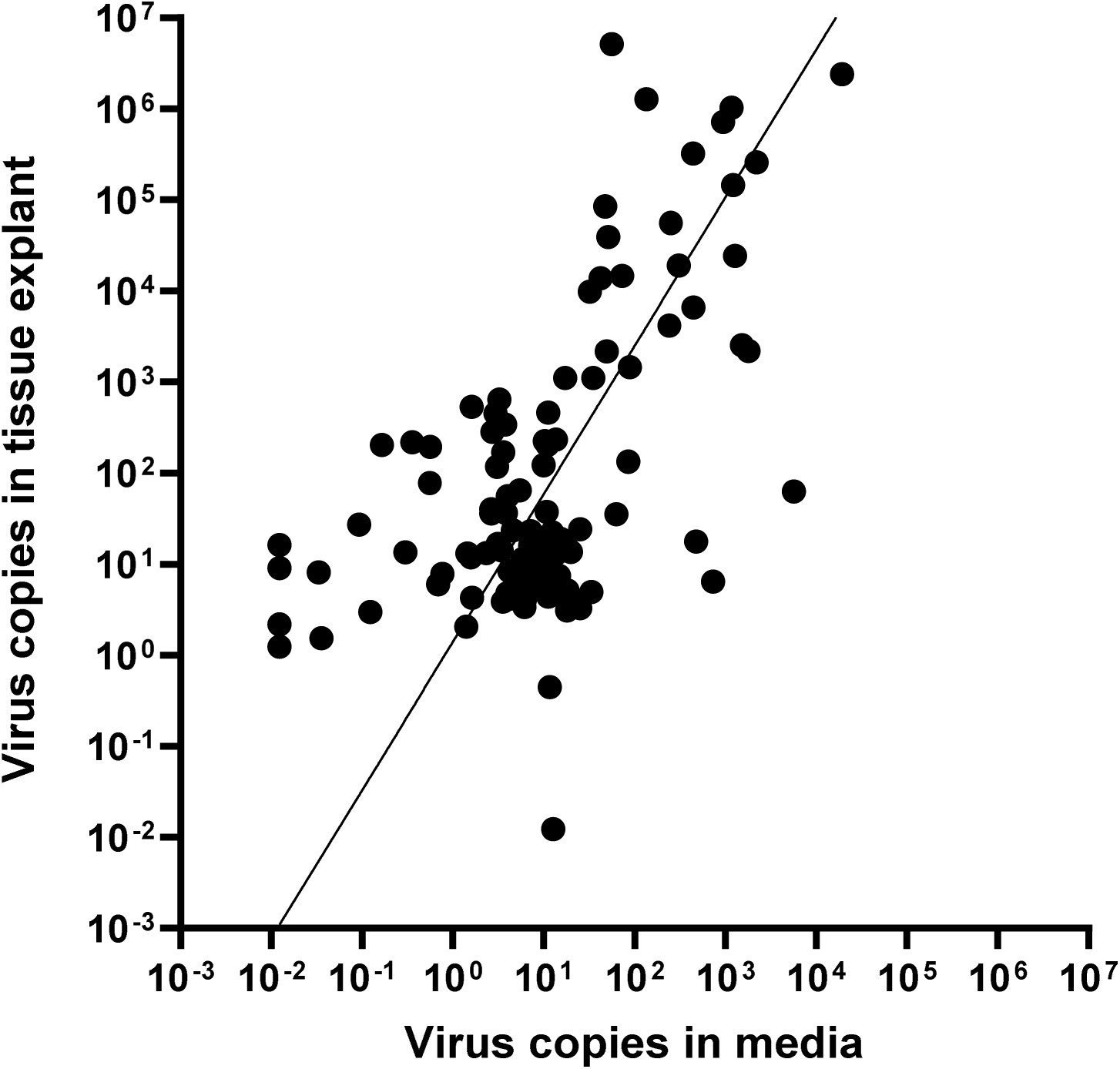
Viral copies in media compared to values from tissue DNA extractions. Non-linear regression fit calculated in GraphPad Y=10^(1.626X + 0.1443). Pearsons R^2^=0.433, p<0.001)

### Histology and Electron Microscopy

The semi-thin electron microscopy (EM) sections showed marginalised chromatin and empty vacuoles, typical of OsHV-1 infected cells. Marginalised chromatin and empty vacuoles were also visible by TEM, as well as fully formed viral capsids (Figure 6A and 6B), similar to available OsHV-1 standards (e.g. Renault et al., 2002). Alongside the molecular data, this is strong evidence that the tissue explant system developed here is capable of replication OsHV- 1 in a controlled environment.

**Figure 6.**
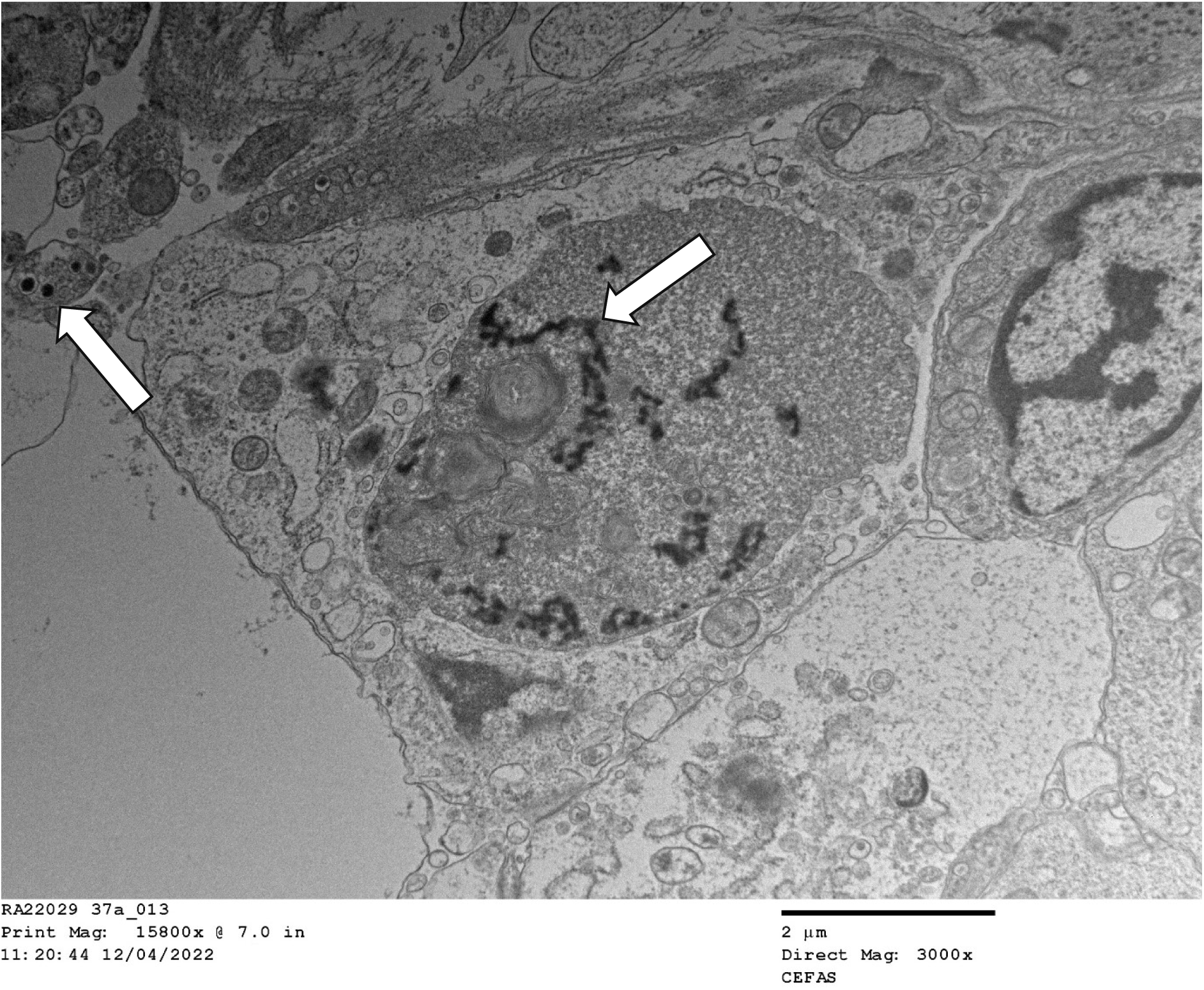
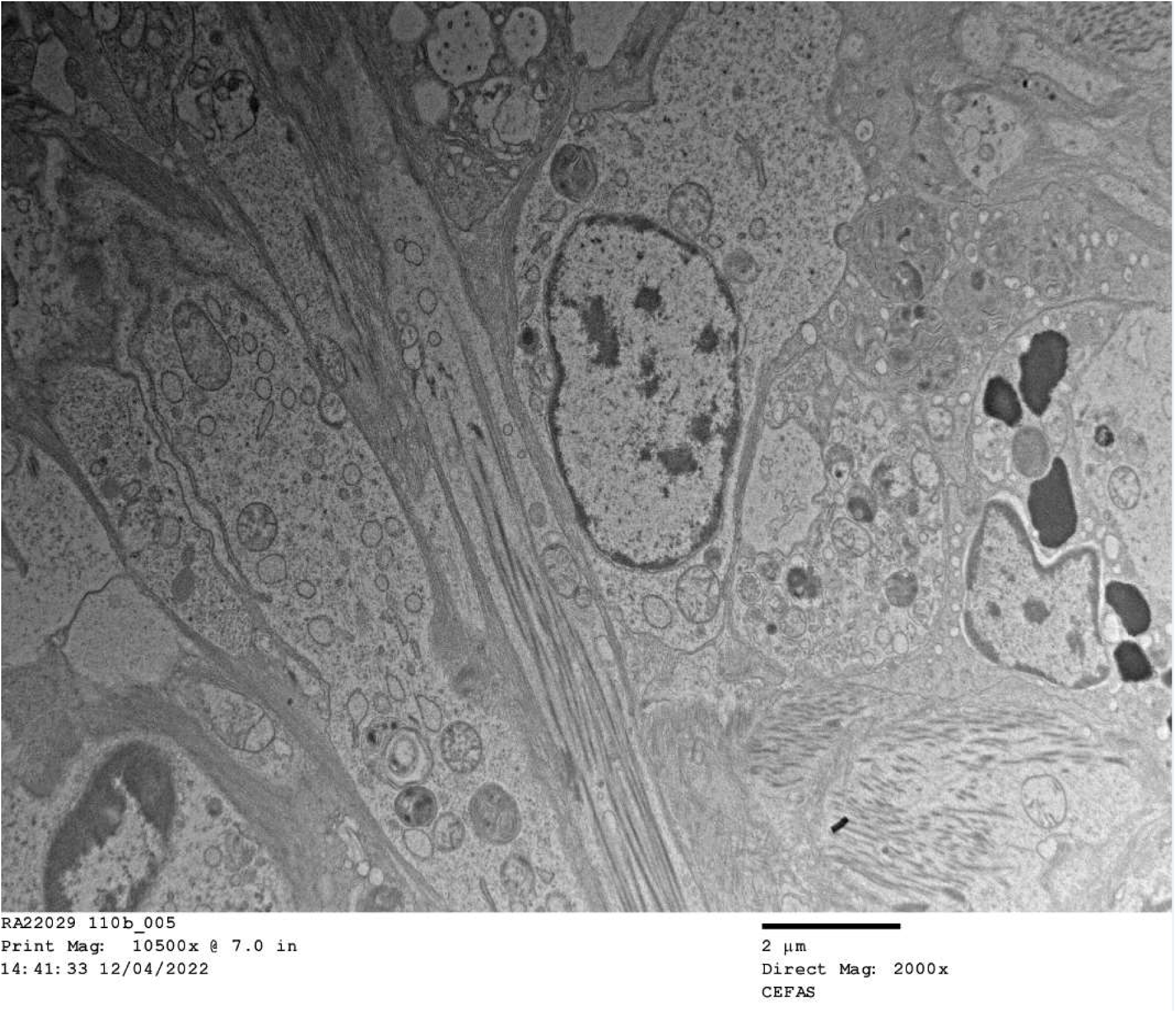
A Electron micrograph of an OsHV infected sample of mantle tissue 6 days post infection harvested 7 days post explant. Arrows show enclosed viral capsids outside nucleus, pre-capsid viral particles inside the nucleus. 6B Electron micrograph of a control sample of uninfected mantle tissue harvested 7 days post explant

## Discussion

This research shows that OsHV-1 can replicate *ex vivo* in tightly controlled axenic laboratory conditions. Samples of explant mantle, muscle and gill tissues all showed clear signs of infection both from molecular and pathological data. Replication of OsHV-1 in controlled conditions opens the possibility to study the dynamics of OsHV-1 infection in isolation, without the influence of other biotic factors such as bacteria and animal behaviour, and abiotic factors such as temperature and salinity changes. This approach may be particularly beneficial for studying the role of genetics in disease resistance without confounding factors. Additionally, the tissue explant infection model removes some of the logistical barriers for OsHV-1 research. Oysters can be shipped alive internationally, direct access to seawater is not required, and biosecurity is considerably simpler to attain, so this approach could be done anywhere, without requirements for complex aquarium systems. *Ex vivo* models will also help to make experiments consistent and repeatable.

The difference in response to infection between the two sources of oysters in whole animal challenge appears to be reflected in the response of the tissue to infection, which was the only variable that significantly explained variation between groups. This suggests that the whole tissue explant model is relevant to aquaculture environments. The system underlying this difference in susceptibility to OsHV-1 is unknown, but this method would allow more detailed analysis of underlying mechanisms. Additionally, this finding helps to further validate the usefulness of the large explant method developed previously (Potts et al 2020), helping to expand on the range and efficacy of tools available for studying OsHV-1 disease dynamics and marine bivalves generally. However, there was a high level of variability within groups from each location, with some tissues becoming infected and others not. It is unclear yet whether this relates to the infection dynamics, which often results in susceptible oysters surviving OsHV-1 exposure (e.g. Figure 3), or inefficiencies in the infection model used.

The whole tissue explant infection model could also potentially help to elucidate the key site of infection within the oyster. Mantle, muscle and gill tissues were all shown to replicate OsHV- 1 *in vitro* and showed different patterns of viral replication, with gill and mantle from site 1 (the more sensitive site) showing significantly higher levels of infection than muscle suggesting they may be more susceptible to initial infection. Hemocytes have been the focal cell type in OsHV-1 pathology (Tracy et al., 2020, Ji et al., 2017, Allam and Raftos, 2015), but as they are found throughout the oyster and in all the tissues examined in this study, it is difficult to confirm that hemocytes are essential in the OsHV-1 lifecycle or if they are infected as a consequence of infection elsewhere in the oyster. Herpes simplex virus (HSV) infects humans and can remain dormant in nerve tissue (Steiner and Kennedy, 1995). Ganglia can be extracted from Pacific oysters, so it may be possible using explants to examine the role of nervous tissue in the OsHV-1 lifecycle (Réalis-Doyelle et al., 2021).

Using OsHV-1 isolated from the tissue explants, or by transferring explants into an environment with naïve oysters and monitoring mortality, morbidity and molecular evidence of infection, would be a useful method to confirm the infectious capabilities of the OsHV-1 produced in explants. This could potentially be used to further examine the dynamics of the infection, including differences between viral replication and shedding in different tissues, from different sources of oyster, or at different times post infection, all studied in isolation with reduced noise compared with conventional in vivo infection models.

An underlying issue with all available models of OsHV-1 infection in oysters is that there is often no way of testing if the oysters used for the challenge have previously been exposed to OsHV-1. Unreported OsHV-1 exposure could lead to inadvertently using resistant animals from within a population, or potentially immune priming of animals making them more resistant to future exposure, thus altering the outcome of experimental disease challenges (Green and Speck, 2018). It is unclear whether using explants instead of whole animals would help to alleviate this issue. Unreported pathogen exposure may be prevalent across marine molluscs where they are grown in an uncontrolled environment, and the extent to which this plays a role in influencing the results of experimental challenges and industrial breeding programmes remains unclear. Populations from both sources of oysters studied here seemed to have some level of survival in OsHV-1 challenges, which could be explained by one of these factors, although the current infection status in the UK suggests both source locations are free from OsHV-1. It is currently impossible to tell whether an oyster is completely naïve to OsHV-1 as their immune system is not well characterised for use of antibody detection, and can be primed by exposure to a different pathogen or physical damage (Melillo et al., 2018, Lafont et al., 2017).

This research also demonstrated the persistence of OsHV-1 DNA in water over time, showing that viral material can be detected in seawater up to 8 days after being present for the first time. Interestingly, OsHV-1 DNA degraded less in ASW, RO water and oyster cell culture media than in natural seawater. The reason for this is unclear, but may be relevant to OsHV-1 monitoring in aquaculture environments. The viability of the virus over time however, appeared to decay rapidly, and after just 48 hours incubation in artificial seawater it was not able to cause any mortality in oysters. The threshold for virulence has not been thoroughly tested, and is known to vary, so it is possible that there is viable virus present but the amount present was below the critical threshold for this particular challenge system. The reservoir of OsHV-1 during periods without outbreaks has not been identified, but this data suggests it is unlikely to be free viable virus within salt water systems (Whittington et al., 2018).

## Conclusion

Using the whole tissue culture system described here, we have shown for the first time that OsHV-1 can replicate *ex vivo* under controlled laboratory conditions. By using two different sources of Pacific oysters from the UK, quantitative PCR revealed that the phenotype of the donor population is reflected in the outcome of infection for the tissue explants, which supports the model’s relevance to the aquaculture environment, and further validates its use for studying the complexities of Pacific oyster diseases. Improvements in the experimental use and understanding of OsHV-1 have also been described, including the rate of decay of virus in seawater compared to other media. These advances should help to focus research on OsHV-1 by providing new systems for study.

## Acknowledgements

The authors acknowledge funding from the Biotechnology and Biological Sciences Research Council (BBSRC), including Institute Strategic Programme grants (BBS/E/D/20002172, BBS/E/D/30002275 and BBS/E/D/10002070), PhD studentship funded within the EastBio DTP programme (BB/M010996/1) and Centre for Environment, Fisheries and Aquaculture Science (Cefas) Seedcorn project DP901W. The authors would also like to thank Kokie Harris and Dr Stefano Carboni for discussions around viral viability experiments.

